# Two residues reprogram immunity receptor kinases to signal in nitrogen-fixing symbiosis

**DOI:** 10.1101/2024.08.22.609144

**Authors:** Magdalini Tsitsikli, Bine W. Simonsen, Thi-Bich Luu, Maria M. Larsen, Camilla G. Andersen, Kira Gysel, Damiano Lironi, Christina Krönauer, Henriette Rübsam, Simon B. Hansen, René Bærentsen, Jesper Lundsgaard Wulff, Sarah Holt Johansen, Gülendam Sezer Kaya, Jens Stougaard, Kasper Røjkjær Andersen, Simona Radutoiu

**Author notes:** Corresponding authors. Kasper Røjkjær Andersen, Simona Radutoiu. Joint first authorship (Magdalini Tsitsikli, Bine W. Simonsen, Thi-Bich Luu).

## Abstract

Receptor signalling determines cellular responses and is crucial for defining specific biological outcomes. In legume root cells, highly similar and structurally conserved chitin and Nod factor receptor kinases activate immune or symbiotic pathways, respectively, upon perception of chitinous ligands. Here, we show that specific amino acid residues in the intracellular part of the Nod factor receptor NFR1 determine signalling specificity and enable the distinction between immune and symbiotic responses. Functional investigation of CERK6, NFR1 and receptor variants hereof revealed a conserved motif that we term *Symbiosis Determinant 1* in the juxtamembrane region of the kinase domain that is key for symbiotic signalling. We demonstrate that two residues in *Symbiosis Determinant 1* are indispensable hallmarks for NFR1-type receptors and are sufficient to convert *Lotus* CERK6 and barley RLK4 kinase outputs to enable symbiotic signalling in *Lotus japonicus*.

## Main

Receptor kinases initiate and regulate diverse signalling events in eukaryotic cells in response to extracellular signals^1,2^. Evolution has diversified the extracellular, ligand-binding domains of these receptors while retaining the coupling to intracellular kinases to maintain the catalytic function^3-5^. Kinases share structurally similar domains but differ greatly in their determinants of specificity mediating the activation of specific downstream signalling pathways^6,7^. Identification and prediction of determinants of signalling specificity has proven a challenge. In plants, Chitin Elicitor Receptor Kinases (CERK) recognize chitin molecules and initiate an immune response^8-10^, and facilitate symbiosis with arbuscular mycorrhizal fungi^11-16^. Legume plants have evolved an additional, but highly similar NFR1 receptor to perceive decorated chitin ligands called Nod factors, produced by symbiotic bacteria to initiate nitrogen-fixing symbiosis^17,18^. Studies based on receptor mutants determined that in legumes, the chitin-triggered immunity and nitrogen-fixing symbiosis are genetically and functionally separated^8,12^ even though both receptors can be expressed in the same cell^19^ (Fig. 1a). This suggests that the receptors themselves encode signalling specificity. Therefore, LysM receptor kinases offer an attractive test system for identifying determinants of specificity to understand how similar kinases control divergent signalling pathways.

**Figure 1.**
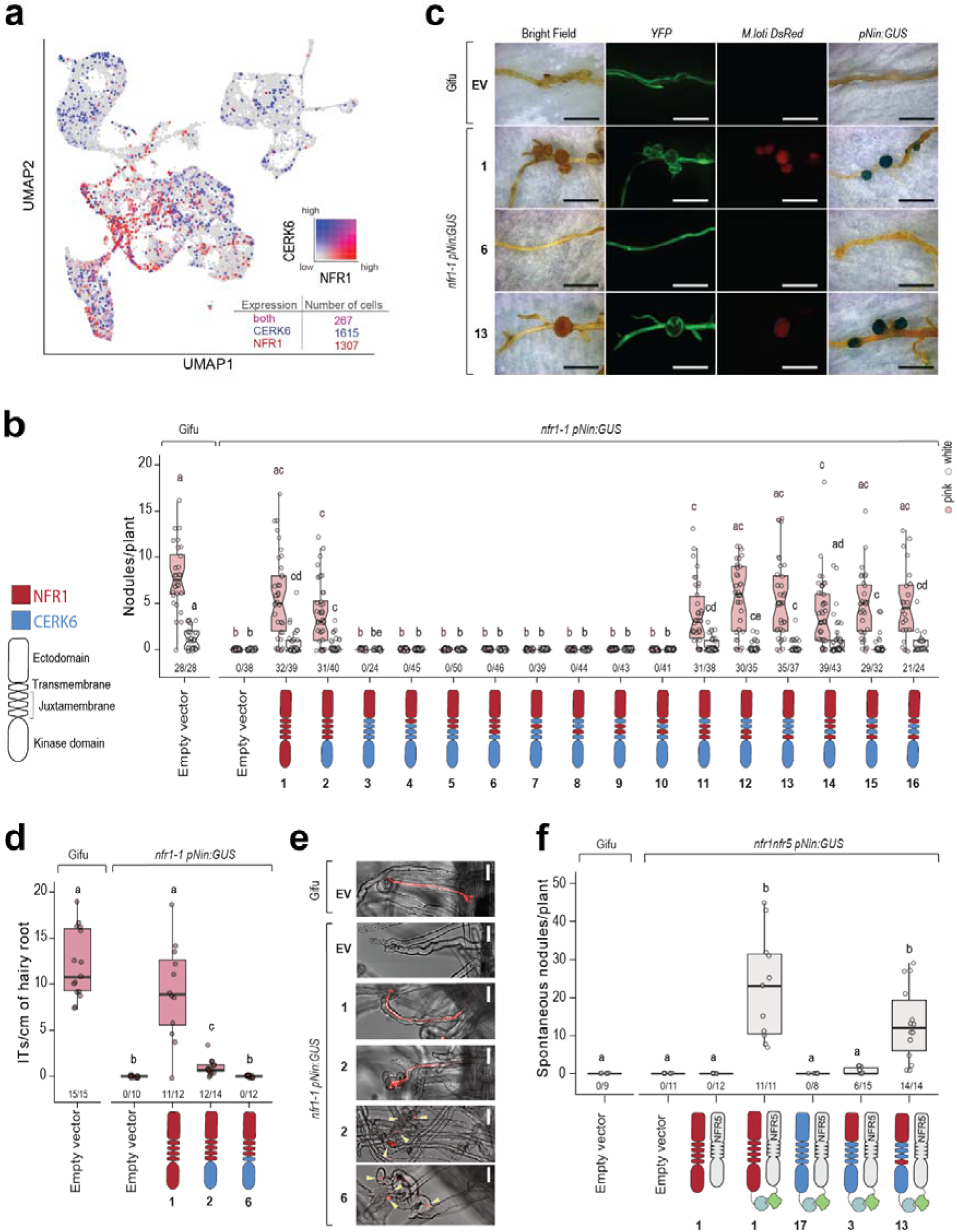
Symbiosis Determinant 1 controls symbiosis with rhizobia. **a** *Lotus* root cells expressing *NFR1* (red), *CERK6* (blue), or both (magenta) 10 dpi (UMAP generated by https://lotussinglecell.shinyapps.io/lotus_japonicus_single-cell/). **b** Number of pink or white nodules^41^ formed by *nfr1* roots expressing the indicated receptor variants. Nodules were counted 5 weeks post inoculation with *M. loti DsRed*. Fractions under the boxplots indicate the number of nodulating plants out of total plants tested. **c** Representative photos of transgenic roots carrying the indicated receptor variants. *YFP*: Yellow Fluorescent Protein (transformation marker), *M. loti DsRed* (fluorescent bacteria), *pNIN:GUS*: Bright Field photos after GUS staining. Scale bars: 5 mm. **d** Number of infection threads (ITs) formed per cm of transgenic root expressing the indicated receptor variant. ITs counted 3 weeks post inoculation with *M. loti MAFF*. ‘*n’* indicates the number of analysed plants. **e** Photos of transgenic roots expressing the indicated receptors at 3 weeks post inoculation with *M. loti MAFF*. Red shows *M. loti MAFF* bacteria. Yellow arrowheads indicate branched and scrambled root hair tips where microcolonies of bacteria are visible. Scale bars: 30 μM. **f** Number of spontaneous nodules formed on *nfr1* roots expressing the indicated constructs. Fractions under the boxplots indicate the number of nodulating plants out of total plants tested. In **b, d** and **f** lowercase letters indicate significant differences based on Kruskal-Wallis analysis of variance and post-hoc analysis (Dunn’s test) p < 0.05.

Previous studies of CERK6 and NFR1 from *Lotus japonicus* (*Lotus*), and LYK3 (the NFR1 homologue) from *Medicago truncatula* (*Medicago*) identified that their ligand affinity and signalling ability depend on distinct regions present in their ectodomains distinguishing chitin and Nod factors^5^. These studies also found that while the ectodomains are in place to recognize the correct extracellular signal, the nature of the cellular pathways is dependent on residues present in the intracellular regions of NFR1 and CERK6 receptors. The CERK6 kinase domain can activate symbiosis only in the presence of the transmembrane-juxtamembrane (TM-JM) region from NFR1. The NFR1 kinase, on the other hand, is not able to activate immune related ROS production after chitin exposure irrespective of the origin of the TM-JM region^5^. This indicates that root nodule symbiosis is controlled independently by specific residues present in the TM-JM region or in the kinase of NFR1, while chitin immunity is controlled by residues present in the CERK6 kinase. The regions in the intracellular part of NFR1 and CERK6 that control this divergent signal output are currently unknown. To discover the specificity determining regions, we performed a detailed functional investigation of chimeric NFR1-CERK6 receptors. Crystal structures informed the design of receptor variants that were functionally tested in *nfr1*, Nod factor insensitive, or in *cerk6*, chitin insensitive loss-of-function mutant plants by expression from their native promoters. We identified key regions and residues that control signalling specificity and demonstrated that engineering symbiotic functionalities into chitin receptors from legume and non-legume plants is possible.

### Symbiosis Determinant 1 defines the signalling specificity of NFR1

Residues present in the transmembrane and juxtamembrane regions have been identified to play distinct regulatory roles for the core kinases of single-pass receptors^20,21^. We aimed to identify residues controlling the specificity of signalling for nitrogen-fixing symbiosis in NFR1^5^. For this, we created a series of fifteen receptor variants where the NFR1 ectodomain was coupled to the CERK6 core kinase via different configurations of the transmembrane (TM) and juxtamembrane (JM) (Fig. 1b, c, receptors 2-16) and investigated their function *in vivo* compared to NFR1 (receptor 1). The first results confirmed that the CERK6 kinase could initiate signalling for root nodule formation and infection only when preceded by the TM-JM of NFR1 (Fig. 1b, receptor 2), but not by the TM-JM of CERK6 (Fig. 1b, receptor 3)^5^. A region spanning 23 residues located in the C-terminal part of the JM adjacent to the kinase domain was found to be critical for the symbiotic proficiency of these receptors (Fig. 1b, c). All variants with the NFR1 version of this region (receptors 11-16) induced formation of infected root nodules expressing the symbiotic marker *Nin*^22^(Fig. 1b, c, Fig. S1). By contrast, all receptors with the corresponding version of CERK6 (receptors 3-10) were unable to induce nodule organogenesis or *Nin* expression in *nfr1* roots, irrespective of the origin of the remaining regions of the JM domain (Fig. 1b, c, S1). We named this region of NFR1 the Symbiosis Determinant 1 (SD1). To verify protein integrity, we expressed the receptor variants in tobacco leaves or *Lotus* root protoplasts and found them to be localized at the plasma membrane, indicating that receptors which are symbiotically inactive are expressed and correctly localized (Fig. S2, S3). Next, we investigated the activation of the symbiotic epidermal program characterized by rhizobial infection via root hair deformation, microcolony and infection thread formation, as well as the early activation of *Nin* promoter in the epidermis. We inspected the root hairs of *nfr1* expressing key receptor variants (receptors 1, 2, and 6) analysed for nodule organogenesis (Fig. 1b). Roots expressing the full-length NFR1 (receptor 1) had typical root hair curling, bacteria entrapment, and long, elongated infection threads (Fig. 1d, e, S4). Such elongated infection threads were also identified in roots expressing the symbiotically functional receptor 2 (Fig. 1b, d, e), albeit in a significantly reduced number, while none were detected for the nonfunctional receptor 6 (Fig. 1b, d, e). Importantly, roots expressing receptors 2 and 6 responded to the presence of bacteria with root hair deformation and microcolony formation even when intracellular root hair infection was not enabled (Fig. S4). This shows that receptor 6 which recognizes the Nod factors via the NFR1 ectodomain initiates only root hair deformation responses in the *nfr1* mutant, but that key events leading to symbiosis, such as the *Nin* promoter activation in the epidermis, intracellular infection, and nodule organogenesis, fail to be activated (Fig.1e, S4). The lack of symbiotic functionality was further confirmed in an independent experiment where the synthetic coupling of receptor 3 to NFR5 using the nanobody-mediated receptor complex formation strategy^23^ showed an inability to induce spontaneous nodule formation (Fig. 1f-receptor 3, 17). However, this was reconstituted if the SD1 was present (Figure 1f-receptor 13) demonstrating the key role of this region in activating the symbiotic signalling from a heteromeric complex with NFR5. Together, our studies demonstrate that SD1 controls root nodule symbiosis ensuring activation of both organogenesis and root hair infection downstream of receptor complex formation.

### The region containing SD1 is a surface exposed motif in the kinase domain

The region containing SD1 has an amino acid composition that distinctly separates Nod factor receptors from their chitin-binding counterparts and other members of this protein family in legume plants (Fig. S5 and Supporting Material 1). To understand the molecular details of SD1, we expressed and purified the intracellular domains of CERK6 and NFR1 (Fig. S6) for structural investigation. The purified proteins were functional and able to bind ATP (Fig. S7). We obtained well-diffracting crystals for CERK6 when an inhibitory mutation, D460N, in the DFG motif was introduced, but this was not the case for the tested NFR1 versions. To obtain crystals of a symbiotic kinase, we turned to the NFR1 orthologue from *Medicago*, LYK3, which crystalized with the same inhibitory mutation, D459N, as CERK6 (Fig. S8). The atomic structures of CERK6 and LYK3 kinases were determined to 1.85 Å and 2.5 Å resolution, respectively (Table S1). Both LYK3 and CERK6 showed a classical fold with clearly defined N- and C-lobes and hallmark structural features of active kinases (Fig. 2a, b). LYK3 and CERK6 are both crystallized in an inactive conformation closely resembling the conformation of the CDK2/Src inactive kinase, which is an autoinhibition state of the kinase, where the αC is in the “out”-position, while the aspartate of the DFG motif points towards the nucleotide binding side. LYK3, but not CERK6, was crystallized with a bound ATP analogue (Fig. S8a). However, despite this, the two structures show an overall high similarity (RMSD=0.724 Å) (Fig. S8b). The SD1 region is clearly defined in the density map of the LYK3 structure as a loop followed by the αB helix, which is a conserved feature of IRAK/Pelle-type kinases. SD1 is kept in place by specific residues that are conserved between chitin and Nod factor receptors, stabilizing the interactions to the N-lobe of the kinase (Fig. 2c). SD1 contains only six residues differing between CERK6 and NFR1, three of these are in the N-terminal loop, while the remaining three are on the αB helix. All six defining residues in SD1 are surface exposed making this motif accessible for interactions (Fig. 2d, e).

**Figure 2.**
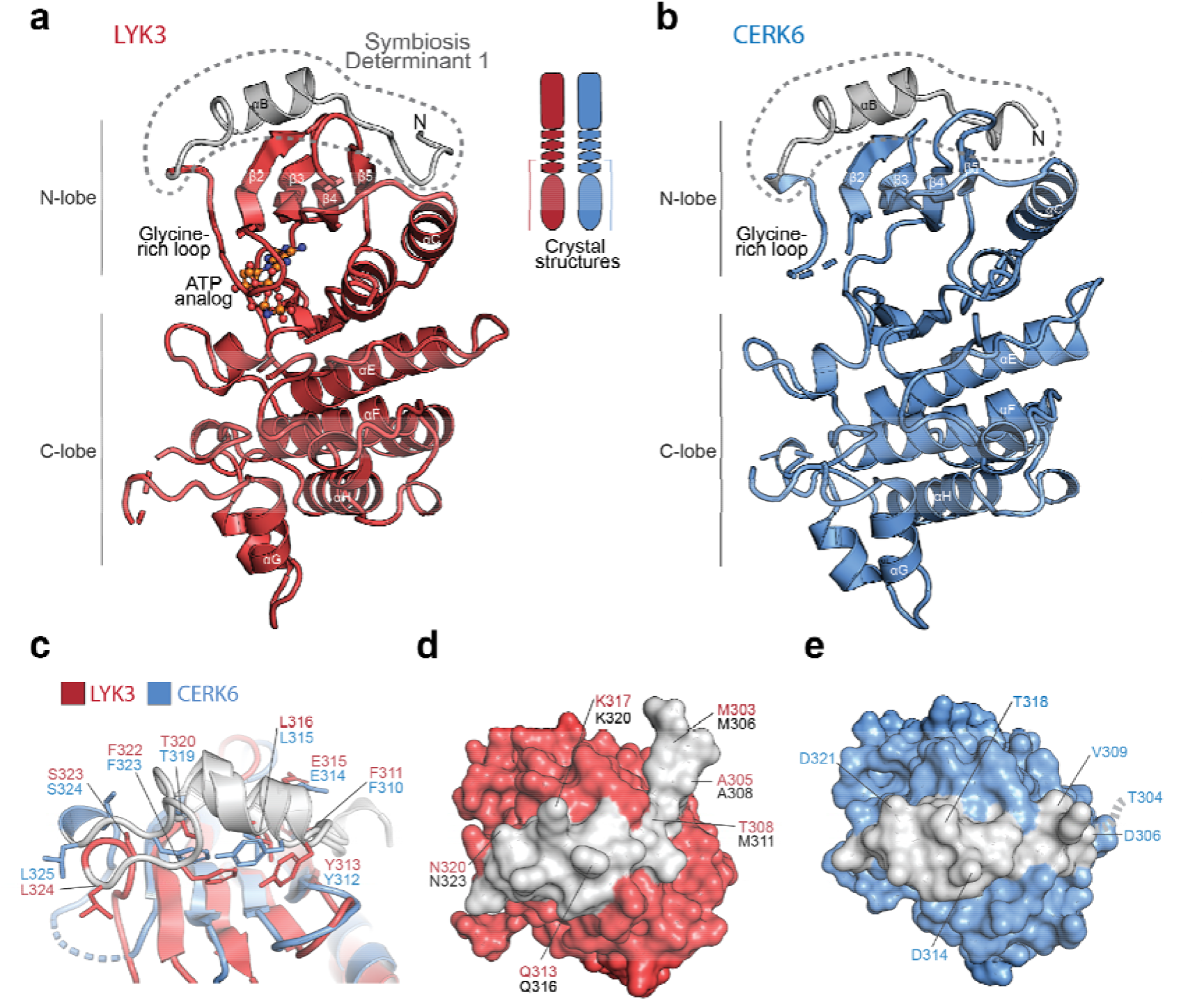
Crystal structures of LYK3 and CERK6 kinase domains. Structure of the LYK3 **(a)** and CERK6 **(b)** kinase domains annotated with conserved secondary structural elements (LYK3, PDB-ID: 9GFZ, CERK6 PDB-ID: 9GB9). LYK3 kinase was crystallized with AMP-PNP in the nucleotide binding site. The identified SD1 region is indicated in gray. **c** Structural overlay of the LYK3 and CERK6 showing the region with SD1. The indicated amino acids that interact with the kinase core domain and SD1 are conserved. **d**-**e** Zoom-in on SD1 of LYK3 and the same region in CERK6 seen from the top showing that this region is surface exposed and accessible. In CERK6, the T304 (dotted line in **e**) is not visible in the structure. In **(d)** residues in red are from LYK3 and those in black are from NFR1.

### The core kinase domains are important for the two diverging signalling pathway

Using the available structural information for the NFR1 and CERK6 receptors, we next aimed to identify which regions or residues in the core kinases control the specificity of the two signalling pathways. Structural overlay revealed that the differing residues between CERK6 and NFR1 were dispersed across both the N- and C-lobes and were primarily localized on the surface (Fig. S9), likely to facilitate protein-protein interactions^6^. Knowing that core kinases of NFR1 and CERK6 can initiate symbiosis or immunity independently of the TM-JM regions^5^, underscores the importance of diverging, surface-localized residues contribute to signalling specificity. To identify contributing surfaces, we investigated which residues of the NFR1 core kinase differing from CERK6 can drive symbiosis independent of SD1. We identified seven regions (A to G) containing multiple different divergent residues that were structurally clustered, defining potential interaction surfaces (Fig. S9, S10). These were exchanged individually from CERK6 to NFR1 in receptor 6, which was symbiotically nonfunctional (Fig. 1b, Fig. S10). None of the surfaces investigated alone was able to activate root nodule formation in *nfr1*, indicating a possible collective contribution (Fig. S10). We next tested if their combined differences might be a limiting factor for symbiosis establishment (Fig. 3a). Receptor 20, where both the αC helix (region E) and the activation loop (region A) of CERK6 were exchanged to those of NFR1, was indeed found to be able to activate the root hair symbiotic program (Fig. S11) and to induce nodule formation on *nfr1* roots, albeit with very low efficiency, when compared to functional receptors 1, and 18 containing the full-length NFR1, or the entire NFR1 kinase (Fig. 3a). This provides further indications that multiple surfaces of the core NFR1 kinase contribute to symbiosis signalling. Therefore, we tested two variants where the N or C-terminal regions of CERK6 were exchanged to NFR1 in receptor 6 (Fig. 3a receptors 21,22, Fig. S11) without a negative impact on protein folding and ATP binding activities (Fig. S5). Receptor 22 which has the N-terminal region of CERK6 and the C-terminal of NFR1 ensured the activation of both epidermal symbiotic signalling (Fig. S11) and formation of functional nodules more efficiently than receptor 20 where only the αC helix and the activation loop of CERK6 have been exchanged to NFR1 (Fig. 3a). The opposite variant, receptor 21 (containing the N-terminal region of NFR1 and the C-terminal of CERK6), did not have the same effect. The symbiotically functional receptor 22 contains 45 residues diverging from CERK6, scattered across several surface areas (Fig. S9, 10). This shows that residues from the C-terminal region of NFR1 create collectively a larger or a particularly well-defined surface enabling symbiosis.

**Figure 3.**
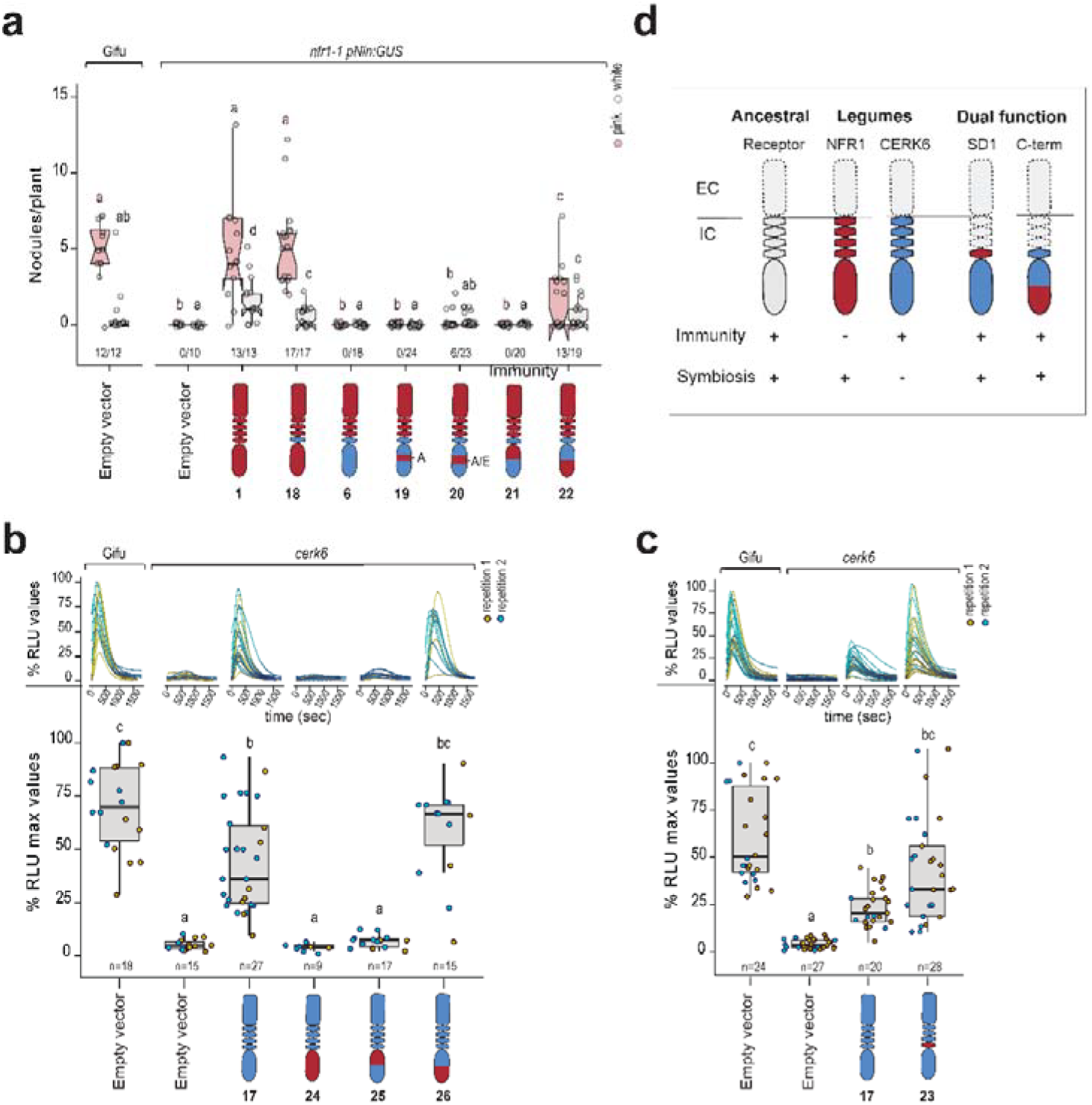
The kinase N- and C-lobes contain determinants controlling independently the two signalling pathways. **a** The number of pink or white nodules formed on *nfr1* roots expressing the indicated receptor variants. Nodules were counted 5 weeks post inoculation with *M. loti DsRed*. Fractions under the boxplots indicate the number of nodulating plants out of total plants tested. Lowercase letters indicate significant differences based on Kruskal-Wallis analysis of variance and post-hoc analysis (Dunn’s test) p < 0.05. **b-c** Reactive oxygen species (ROS) generated by *cerk6* roots expressing the indicated receptor variants within a period of 30 min after treatment with 0.1 mg/ml chitin. Top: curves show the values of RLU (Relative Luminescence Units) during the 30-min period after chitin treatment. Bottom: boxplots show the maximum value of RLU observed for each sample within the 30-min period after chitin treatment. (100% is the highest value observed in WT expressing the empty vector construct). **d** Schematic representation of the observed functional capacities for the intracellular domains (IC) of the indicated LysM receptors. These capacities are dependent on the capacity of the ectodomains (EC), to recognize and bind Nod factors or chitin^5^.

The NFR1 core kinase does not have the ability to activate immunity^5^, and our findings show that residues in its C-terminal region have diverged from CERK6 to enable symbiosis (Fig. 3a, d). This raises the question of whether the specificity for symbiotic signalling evolved in the core kinase at the expense of immunity signalling. We investigated this possibility by exchanging the C-terminal region of the full-length CERK6 with the symbiosis-permissive region of NFR1. This receptor version expressed in the *cerk6* mutant from the *Cerk6* promoter was able to induce the production of reactive oxygen species (ROS) after chitin treatment (Fig. 3b, receptor 26), indicating no negative interference from these NFR1-type residues on this signalling pathway. By contrast, receptor 25 containing the NFR1-type residues in the N-terminus of the kinase failed to complement the mutant phenotype (Fig. 3b, receptor 25) indicating that the N-terminal region of CERK6 contains determinants for immune signalling. The C-terminal region of the NFR1 core kinase acts independently of SD1 in activating symbiosis (Fig. 3a, receptor 22), and importantly, it does not interfere in immunity. Therefore, we asked if the SD1 in NFR1 may affect the immune signalling from the CERK6 core kinase. We exchanged the corresponding region with the NFR1 variant within the full-length CERK6 (Fig. 3c, receptor 23) and found that this receptor variant 23 also induced production of ROS after chitin treatment when expressed in *cerk6* mutant. Together, these results demonstrate that neither of the two regions found here to be determinants for symbiosis have a negative impact on the capacity of CERK6 to activate ROS and that the functionality from the intracellular region of CERK6 or NFR1 can be independently modulated by residues located in distinct regions (Fig.3d, e).

### Specific residues in SD1 coordinate organogenesis and infection

All land plants contain CERK receptors that activate immunity, but only legumes contain NFR1-type receptors that activate root nodule symbiosis^24-26^. The identification of two distinct regions in NFR1 extending the molecular functions of CERK6 prompted us to investigate if this can be employed for engineering of CERKs. The C-terminus of the NFR1 core kinase contains 45 residues diverging from CERK6 and our extensive analysis of receptor variants revealed a synergetic action of these variable residues (Fig. 3a). On the other hand, the SD1 contains only 6 residues that differ from CERK6 (Fig. 4a, Fig. S5) and are thus more amenable to engineering. Therefore, we asked if all defining residues of SD1 are crucial for the symbiotic program. We examined variants of the symbiotically active receptor 2 where individual variable residues of SD1 were exchanged from NFR1 to CERK6. (Fig. 1b, 4b). All these variants retained the ability to induce nodule formation after inoculation with *M. loti*, indicating a synergetic contribution to signalling. Closer inspection of the phenotypes, however, revealed residue-specific variations (Fig. 4b, c, Fig. S12, S13). Receptor 27 (M306T) ensured the formation of both infected and white nodules albeit on fewer plants (18 out of 24) and in a significantly reduced number compared to receptor 2 (29 out of 30) (Fig. 4b, c). This indicates an impact of M306 on the efficiency of symbiotic signalling. Receptor 28 (A308D) had the lowest symbiotic capacity. Only 15 of the 40 *nfr1* roots expressing this receptor formed infected nodules, a significantly reduced number when compared to those expressing receptor 2 (Fig. 4b, c, Fig. S12). This shows that A308 in SD1 is critical for root nodule formation. Nodule organogenesis and rhizobial infection are well coordinated processes in symbiosis and the phenotypes observed in receptors 27 and 28 imply that variable residues of SD1 may not only control the initiation of symbiotic signalling, but they may also coordinate organogenesis and infection. To investigate this, we inspected the transgenic roots in more detail for the expression pattern of the *Nin* promoter providing a better insight on the early signalling events. No infection threads or epidermal *Nin* activation were detected at two weeks post inoculation when receptors 27 or 28 were inspected for early symbiotic events (Fig. S13). At 5 weeks, however we found that *nfr1* roots expressing receptors 28 (A308D), 30 (Q316D) and 31 (K320T) showed activation of the cortical program where a significantly large number of nodule primordia were present inside the roots, and only a few were developed into nodules. These primordia were only detectable after monitoring for cortical expression of the *Nin* promoter (Fig. S12). It is well-established in root nodule symbiosis that numerous primordia are initiated if there is an impairment of infection^27-30^. Thus, the large number of *Nin*-expressing organogenetic events initiated by these specific receptors indicates that signalling from NFR1 via SD1 is key for the efficiency of signalling as well as for coordinating organogenesis and infection programs.

**Figure 4.**
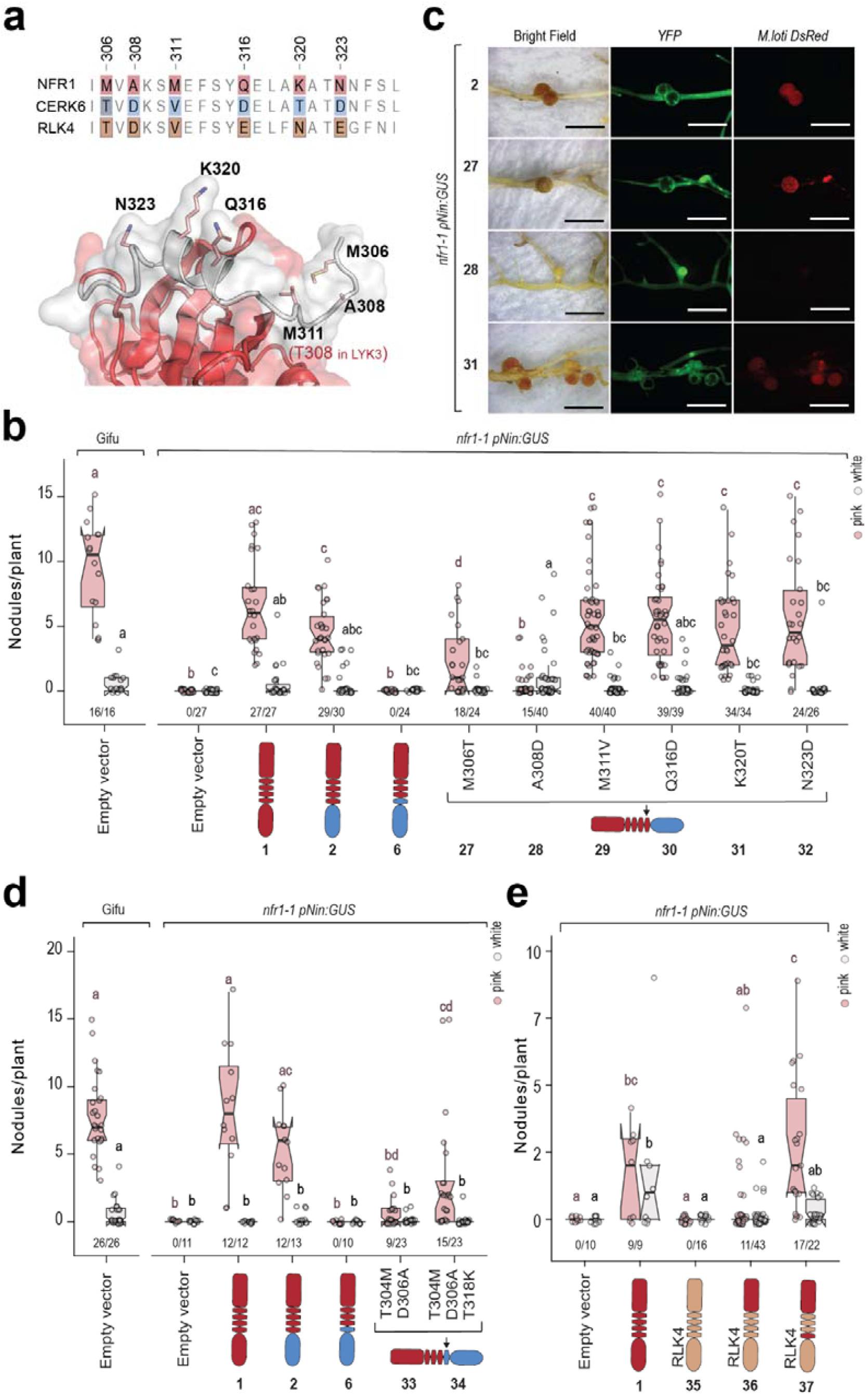
Two residues of Symbiosis Determinant 1 are essential for root nodule symbiosis. **a** Amino acid alignment of Symbiosis Determinant 1 (SD1) in NFR1, CERK6 and RLK4. Numbers above the alignment indicate the amino acid positions in NFR1 protein. **c** Representative photos of transgenic roots expressing the indicated receptor variants. YFP: Yellow Fluorescent Protein (transformation marker), *M. loti DsRED* (fluorescent bacteria), *pNIN:GUS*: Bright Field photos after GUS staining. Scale bars: 5 mm. **b, d**, and **e** Number of pink or white nodules formed on *nfr1* roots expressing the indicated receptor variants. Nodules were counted 5 weeks post inoculation with *M. loti DsRed*. Fractions under the boxplots indicate the number of nodulating plants out of total plants tested. Lowercase letters indicate significant differences based on Kruskal-Wallis analysis of variance and post-hoc analysis (Dunn’s test) p < 0.05.

### Two residues reprogram chitin receptors to signal in symbiosis

Knowing that a synergetic action of the residues from SD1 is needed, we asked whether minimal engineering could be achieved to reroute the signalling from the intracellular region of CERK6 for root nodule symbiosis. As proof-of-concept we targeted the symbiotically nonfunctional receptor 6, where the SD1 and kinase domain derived from CERK6 (Fig. 1b, c). Changing individual residues in the SD1 of CERK6 did not reconstitute symbiosis of the *nfr1* mutant (Fig. S14), but T304M and D306A substitutions together were sufficient to complement nodule formation and rhizobial infection (Fig. 4d, receptor 33) demonstrating that these two residues (methionine and alanine) are the main drivers of specificity for symbiotic signalling. One additional change, T318K, significantly improved the efficiency of the symbiotic signalling (Fig. 4d, receptor 34). Next, we attempted to engineer RLK4, the closest receptor of NFR1 and CERK6 in barley^23^. The kinase of RLK4 is, like CERK6, able to induce immunity by producing ROS in *cerk6* upon chitin treatment (Fig. S15), but unlike CERK6 it has a very limited symbiotic signalling capacity when coupled to the ectodomain of NFR1. Eleven out of forty-three *nfr1* roots expressing receptor 36 with the intracellular region of RLK4 induced a barely detectable symbiotic activity in the roots and produced only one or two nodules (Fig. 4e, S16, S17, receptor 36). This is different compared to CERK6 (Fig. 1b-receptor 3). Nonetheless, the residues within the region of SD1 are conserved between CERK6 and RLK4 (Fig. 4a). Therefore, we asked if engineering this region in the cereal receptor would have a similar impact on symbiosis signalling. We created an engineered receptor 37 where the intracellular RLK4 contained SD1. This receptor, like receptor 13 (SD1 engineered in CERK6), was also able to efficiently initiate root nodule formation and rhizobial infection when expressed in the *nfr1* mutant (Fig. 4e, S16, S17 receptor 37). Unlike CERK6, engineering SD1 in the intracellular region of RLK4 abolishes its capacity to activate ROS production (Fig. S15). Together, these results demonstrate that SD1 provides both *Lotus* CERK6 and barley RLK4 with symbiotic signalling capacity leading to root nodule organogenesis and intracellular bacterial infection.

## Discussion

The ability to detect specific signals present on the cell surface and to respond with the appropriate internal signalling pathway is important for all life forms. Receptor kinases play a major role in eukaryotes in relaying the correct information in the cell, and despite high structural conservation^2,6,31-33^, little is known about how signalling specificity is ensured in plants. Here we investigated the mechanism enabling NFR1 and CERK6 to initiate symbiosis or immune signalling pathways. NFRs evolved as homologs of CERKs and became neofunctionalized in legumes to enable root nodule organogenesis and intracellular infection of nitrogen-fixing bacteria^18,25,34,35^ (Fig. 3d). When Nod factor recognition was ensured by the NFR1 ectodomain, the intracellular region of CERK6 was able to activate root hair deformation, albeit in a rather uncontrolled manner (Figure 1e-receptors 2 and 6), a phenotype previously observed for the symbiotic *nin* mutants^22^. Thus, it is conceivable that specific residues have diverged from CERK receptors to enable initiation of the epidermal infection and cortical root nodule organogenesis. We demonstrate here that two regions, the SD1 in the juxtamembrane and the C-terminus of the NFR1 core kinase, contain residues that control the activation of *Nin* and determine symbiosis signalling. These regions can activate symbiosis independently since the necessity for SD1 can be overcome by NFR1 residues present in the C-terminal region of the kinase and vice versa (Fig. 1b and Fig. 3a, d). Interestingly, an intracellular conformation where the C-terminus is of the NFR1-type can serve a dual function, initiating both symbiosis and immunity, depending on the type of the ectodomain (Fig. 3a, b). This is reminiscent to a certain extent of the functionality of barley RLK4 which was able, albeit with a very low efficiency, to induce root nodule formation (Figure 4e). LYK receptors of *Parasponia andersonii* (Parasponia)^25^ and those from plants of the FaFaCuRo clade forming nitrogen fixing symbiosis with Nod factor-producing Frankia^36^ also lack the SD1 (Figure S4). The LYK1, LYK3a and LYK3b from Parasponia were shown to function in both symbiosis and chitin immunity^25^. SD1 is thus a specific signature of Nod factor receptors from legumes, and we show here that the motif is necessary and sufficient (Fig. 4) for the initiation and coordination of root nodule organogenesis and root hair infection which in legumes runs efficiently. It is thus conceivable that the evolution and maintenance of SD1 in the legume clade have contributed to the evolution of a very efficient symbiosis in Papilionoids^37^.

The residues in SD1 are surface exposed (Fig. 2d, e), allowing this motif to be accessible for potential interactions. Detailed inspection of CERK6 and LYK3 crystal structures revealed no evidence of intra-molecular interactions between variable residues and known regulatory regions of the core kinase that could explain a direct control over signalling (Fig. 2). However, kinases have a dynamic structure^33,38,39^ and both intra- and intermolecular interactions between protomers or with interacting partners e.g. NFR5 may occur once the compatible ligand is bound by the ectodomains. Deciphering the specific contribution of SD1 in NFR1 remains a future challenge due to the inherent structural plasticity of the juxtamembrane region that includes SD1, the temporally dynamic landscape of their interacting partners, and the fact that SD1, specifically regulates the signalling output of the NFR1-NFR5 heteromeric complex (Fig. 1f). The necessity of a large surface in the C-terminal region of NFR1 kinase corroborates with a scenario whereby multiple surface-exposed residues contribute to symbiotic signalling possibly by creating more stable protein-protein interaction interfaces or by enabling dynamic interactions with multiple partners.

Minimal changes were identified here that enabled chitin receptors, CERK6 from *Lotus* and RLK4 from barley, to activate the symbiotic program by inducing expression of *Nin* that controls root hair infection and nodule organogenesis in the cortex. Changing two residues, T304M and D306A, was sufficient for the CERK6 kinase to induce the formation of nodules (Fig. 4d). The symbiotic proficiency was improved by changing T318K and NFR1-levels of nodulation were reached when the entire SD1 was present (Fig. 1b and Fig. 4d). Importantly, our detailed investigations revealed that engineering the intracellular regions of chitin receptors extends their functional capacities and that both symbiotic and immune functionalities can be controlled from a singular version of the intracellular domain. Together with the opportunity of minimal alterations, we provide the blueprint for engineering native receptors to support root nodule symbiosis in non-legumes^40^.

## Acknowledgments and funding

We would like to thank Lene H. Madsen for expertise on gene cloning and Lotus resources, Majken K. Sørensen for plant care and seed production in our greenhouse, and Taylor G. Fitzgerald for proofreading. MT thanks Thomass Ott for support. This work is funded by the Novo Nordisk Foundation (NNF18OC0052855) and the project Enabling Nutrient Symbioses in Agriculture (ENSA), which is funded by Bill & Melinda Gates Agricultural Innovations (INV-57461), the Bill & Melinda Gates Foundation and the Foreign, Commonwealth and Development Office (INV-55767).

## Competing interests

Aarhus University has filed a provisional patent application authored by SR, KRA, MT, BWS, CS, CK, MML, DL, SBH and KG, on using these findings for engineering LysM receptor kinases. The other authors declare no competing interests.

## Author contributions

Conceptualization: SR, KRA.

Methodology: SR, KRA, KG, TBL, CK, DL, MT, BWS, SH.

Investigation: MT, BWS, TBL, CGA, MML, HR, GS, SH, JLW, SHJ. Visualization: MT, BWS, KRA, KG.

Funding acquisition: SR, KRA. Project administration: SR, KRA.

Supervision: SR, KRA, KG, TBL, CK, DL. Writing – original draft: SR.

Writing – review & editing: MT, BWS, TBL, CGA, MML, KG, KRA.

## Notes

### Competing Interest Statement

A provisional patent application authored by SR, KRA, MT, BWS, CS, CK, MML, DL, SBH, KG, on using these findings for engineering LysM receptor kinases has been filed. The other authors declare no competing interests.

### Summary of Updates

This version contains a revised manuscript text, figures and additional data.

